# Forest loss in Indonesian New Guinea: trends, drivers, and outlook

**DOI:** 10.1101/2021.02.13.431006

**Authors:** David L.A. Gaveau, Lucas Santos, Bruno Locatelli, Mohammad A. Salim, Husnayaen Husnayaen, Erik Meijaard, Charlie Heatubun, Douglas Sheil

## Abstract

The rich forests of Indonesian New Guinea are threatened. We used satellite data to examine annual forest loss, road development and plantation expansion from 2001 to 2019, then developed a model to predict future deforestation in this understudied region. In 2019, 34.29 million hectares (Mha), or 83% of Indonesian New Guinea, supported old-growth forest. Over nineteen years, two percent (0.75 Mha) were cleared: 45% (0.34 Mha) converted to industrial plantations, roads, mine tailings, or other uses near cities; 55% (0.41 Mha) cleared by transient processes including selective natural timber extraction, inland water bodies-related processes, fires, and shifting agriculture. Industrial plantations expanded by 0.23 Mha, with the majority (0.21 Mha; 28% of forest loss) replacing forests and reaching 0.28 Mha in 2019 (97% oil palm; 3% pulpwood). The Trans-Papua Highway, a ~4,000 km national investment project, increased by 1,554 km. Positive correlations between highway and plantations expansion indicate these are linked processes. Plantations and roads grew rapidly after 2011, peaked in 2015/16, and declined thereafter. Indonesian government allocated 2.62 Mha of land for the development of industrial plantations (90% oil palm 10% pulpwood) of which 74% (1.95 Mha) remained forest in 2019. A spatial model predicts that an additional 4.5 Mha of forest could be cleared by 2036 if Indonesian New Guinea follows similar relationships to Indonesian Borneo. We highlight the opportunities for policy reform and the importance of working with indigenous communities, local leaders, and provincial government to protect the biological and cultural richness still embodied in this remarkable region.

## 1. Introduction

New Guinea, the world’s largest tropical island, covers just 0.53% of the Earth’s land (785,753 km^2^), but is the most floristically diverse island in the world (Cámara-Leret et al. 2020). It hosts extensive old growth forests, including extensive mangroves and peat swamps, in near pristine settings (Corlett and Primack 2011; Sasmito et al. 2020). The region’s remarkable biota — such as the birds-of-paradise — have fascinated biologists for centuries, although much of the region’s species remains “astoundingly little-known” (Beehler, 2007). A rapid botanical assessment in the foothills of the Foja Mountains in Indonesian New Guinea distinguished 487 plant morpho-species in 15 small plots and found that 156 (32 %), could not be matched to herbarium specimens or published references suggesting a high proportion of undescribed species (van Heist et al. 2010). With such incomplete assessments, it is likely that many of the important locations for biodiversity conservation have not yet been identified. Similar richness and endemism are found among the fauna. The 552 bird species recorded in Indonesian New Guinea include 25 birds of paradise, and a similar number of kingfishers, parrots, raptors, and pigeons along with three cassowaries (Marshall and Beehler, 2007). As with plants, much remains unknown about this fauna: for example, even bird-of-paradise species are still being discovered (Scholes and Laman 2018). The region’s rich biological diversity is the result of a turbulent geological history and its variety of tropical habitats from sea level, climbing up to mountain ranges reaching 4,884 meters, the highest land area between the Himalaya mountains and the Andes (Kooyman et al. 2019; Lohman et al. 2011; Morley, 2000; Toussaint et al. 2014).

Divided in half by its colonial history, the island comprises the nation of Papua New Guinea in the East with the remaining area of 411,387 km^2^ (53%) in the West, part of Indonesia. Indonesian New Guinea, also known as *Tanah Papua* and formerly as *Irian Jaya* — comprises the provinces of Papua (312,806 km^2^) and West Papua (98,581 km^2^). These provinces suffered fewer human impacts than most tropical regions (Brooks et al., 2002; Supriatna et al. 1999) and remain relatively untouched compared with Western Indonesia (Java, Sumatra and Indonesian Borneo (Kalimantan)). They are home to diverse indigenous human cultures each with their own relationship to nature and natural resources (Boissiere and Purwanto 2007; Kennedy 2012), and until recent decades most human impacts appear to have been relatively localized (Bartlett et al. 2016; Johnson et al. 2016; O’Connell and Allen 2015). This is changing.

Indonesian New Guinea has begun to experience large-scale conversion of forests into plantations (Runtuboi et al. 2020). With diminishing agricultural land available elsewhere in the country, plantation companies (oil palm and pulpwood) are increasingly investing in the development of plantations in Papua and West Papua provinces, while the Indonesian government has invested in ambitious road construction programmes to facilitate land development (Franky and Moragan 2015; Sloan et al. 2019). Road developments are also aimed at delivering healthcare, education, and sustainable economic opportunities to remote areas, but many observers are concerned that such developments will accelerate forest conversion as it did in Sumatra and Indonesian Borneo (Kalimantan) (Austin et al. 2019). Construction of the planned ~4,000 km Trans-Papua Highway, a road started in 1979 to link all the major urban centres of Papua and West Papua provinces (Kirksey and Bilsen 2002), has accelerated after 2000. Construction of another major highway intended to ease the development of a food estate project, the Merauke Food Estate and Energy Project, in the south of the island began in 2012. The Indonesian government degazetted 1.2 million hectares (Mha) of State Forest (*Kawasan Hutan*) in 2012 to facilitate these planned roads, oil palm and food estate developments (Mutia Dewi 2020). In 2016, the president of Indonesia is reported to have allocated 85.7 trillion rupiah (~USD6.4 billion) to fund such development projects (including other infrastructure projects like sea ports) (Gokken 2017).

The provincial governments of Papua and West Papua provinces recognize the risks of the business-as-usual plantation and road-led development strategy pursued previously in Sumatra and Indonesian Borneo. These development models have driven deforestation and negatively impacted indigenous communities (Margono et al. 2014; Santika et al. 2019). In 2018, the Governors of Papua and West Papua provinces signed the Manokwari Declaration that commits them to conserve at least 70% of the region, ensure infrastructure developments are “environmentally appropriate” and based on principles of sustainable development while also protecting the rights and roles of indigenous communities (Cámara-Leret et al. 2019). Other goals included developing the information management and monitoring systems needed to support and oversee sound conservation spatial planning and implementation.

While various studies have examined conservation concerns and land cover change in Papua New Guinea (Alamgir et al. 2019; Filer et al. 2009; Laurance et al. 2011; Shearman and Bryan, 2011; Shearman and Bryan, 2015), Indonesian New Guinea is, by comparison, neglected. When it is considered it is typically part of a broader regional overview without concern for local details (Austin et al. 2019; Carlson et al. 2018; Miettinen et al. 2011). An exception is a recent study highlighting regional road developments (Sloan et al., 2019). Here, we use annual Landsat imagery (spatial resolution: 30 m × 30 m) to estimate forest loss, and road and plantation expansion from 2001 to 2019. We identify the distribution and nature of recent forest cover loss and land developments across the region. We examine temporal correlations between these variables and discuss their patterns. We assess other drivers of loss, such as impacts of logging, mining, and other processes. We overlay these data with detailed concession and protected area maps and develop a spatially explicit model to generate a deforestation risk map and assess possible future scenarios.

## 2. Methods

### 2.1. Forest area

Here we are primarily concerned with old-growth forests (here abbreviated to “forests”). These relatively intact closed-canopy (>90% cover) and high carbon stock (Above Ground carbon: 150-310 Mg C/Ha) forests growing, and regenerating naturally, on dry mineral soils and in swamps. Our definition includes intact as well as selectively logged forests. Intact forests have not been severely disturbed by humans in recent decades, or disturbances were too old to be detected by the satellites. Selectively logged forests include old-growth forests that have been impacted by both artisanal tree cutting as well as by more extensive mechanized timber cutting and extraction. There is variation among all these forests. For example, on peat domes, forests may be thinner, low carbon stock pole forests. In coastal regions, forests include mangroves as well as natural stands of Sago palm (*Metroxylon sagu* Rottb.) (Karim et al. 2008). In the drier regions of southern Papua province, on the island of Pulau Dolok in Merauke regency, forests include a unique tall mixed seasonally inundated savanna forest, with trees reaching up to 40 m in height, and 50-60% canopy cover and usually dominated by trees of the Myrtaceae family, such as *Melaleuca cajeputi* Powell and *Lophostemon suaveolens* (Sol. ex Gaertn.) Peter G.Wilson & J.T.Waterh. (widely known by its older synonym *Tristania suaveolens* Sm.) (Marshall and Beehler 2012). Young forest regrowth, scrublands, tree plantations, agricultural land, and grasslands and non-vegetated areas are excluded. Intact and selectively logged or generally disturbed forests are synonymous with “Primary” and “Secondary” forests on the Indonesian Ministry of Forestry and Environment’s maps (MoEF 2018).

We mapped the extent of old-growth forests circa 2000 using cloud-free Landsat composite imagery (See §2.6). The old-growth forest map was created by supervised classification in a Random Forest classifier into ‘Forest’ and ‘Non-Forest’ classes. We trained the model to extract mangrove forests because these forests have a distinct spectral response and location (coastal) compared with other forest (Giri et al. 2011). We also used a peatland map from Indonesia’s Ministry of Agriculture to identify peat-swamp forests (Ritung et al. 2011). We called remaining forests, ‘other forests’, mainly forests on mineral soils, and in non-peat swamp locations.

### 2.2. Forest Loss

We estimated “annual forest loss” as the area of forest (as defined above) cleared by conversion or heavy degradation each calendar year from 2001 to 2019. This measurement is based on the most recent annual Tree Loss dataset (v.17) developed at the University of Maryland with Landsat time-series imagery (Hansen et al., 2013). The Tree Loss dataset measures the removal of trees (tree height >5m) if the canopy cover of a 30 m × 30 m land unit (one Landsat pixel) falls below 30%. It does not distinguish between removal of natural forests (canopy cover >90%, tall trees) and planted trees (where canopy cover can also be >90%). To reduce any error resulting from this ambiguity, we only included *Tree Loss* pixels inside of areas that our analysis identified as forests.

The measurement of forest loss detects conversion to industrial and smallholder plantations (e.g. oil palm, acacia, coffee, rubber, cacao, nutmeg) plantations, other types of agriculture (e.g. paddy fields), urban expansion (including transmigration sites), roads, and mining activity (e.g. digging for gold, or mine tailings). It can also include clearings in the forest caused by natural timber extraction, forest fires and other transient processes (e.g. movement of surface water). Forest loss in the forest canopy caused by the movement of surface water (canopy damage and death of trees caused by meandering rivers, expanding lakes, river overflow and surface run-offs) is usually temporary but forest regrowth will depend on damage inflicted and rivers changing course. Forest fires (canopy damage, death of trees, burnt understorey) will cause lasting forest loss if fires are sufficiently frequent. A forest that burns multiple times over just a few years is typically replaced by fire-tolerant vegetation, often dominated by grasses and ferns. If the forest is impacted by fire once, the larger trees survive and the forest regenerates quickly, except on peatlands, where one fire event may kill all emergent trees if it burns through the roots. Forest loss caused by cutting and extraction of timber (logging roads and clearings where trees are harvested) is usually temporary, as disused tracks and clearings in the forest interior recover after harvesting. In those cases where forest loss is temporary, the term forest degradation rather than forest loss is commonly used.

### 2.3. Expansion of industrial plantations

We define a plantation as an area of land planted with oil palm trees *(Elaeis guineensis* Jacq.), or with a single species of fast-growing trees, most commonly acacia (*Acacia mangium* Willd., and *Acacia crassicarpa* A.Cunn ex Benth.). An industrial plantation typically covers several thousand hectares of land. It is an area bounded by long straight lines, often forming rectangular boundaries. Harvesting trails are usually developed in straight lines forming a rectilinear grid. Annual expansion of industrial plantations represents the area of land (whether forest or non-forest) that has been converted to industrial plantation of oil palm or acacia each year. To map plantations, we scanned a sequence of annual cloud-free Landsat composites forward in time from 2000 to 2019 (see §2.6). We declared an area “industrial plantation” the moment an area that presented the characteristics of industrial plantations (area bounded by several straight lines; spectral colours changing from green to bare land, dense network of trails forming a rectilinear grid) appeared on our sequence of imagery (Figure A1). We delineated the boundaries of plantations at a scale of 1:50,000 using a visual, expert-based interpretation method. We performed this task in the GIS software ArcMap (10.4.1). We also employed maps of oil-palm and pulpwood (acacia) concessions (see §2.7) to further identify the location of plantations. New plantations can either cause deforestation by directly replacing forests or avoid deforestation by planting on low-to-no-tree-covered landscapes. We determined the area of forest (and non-forest) converted every year into industrial plantations. This was ascertained by measuring the overlap between the annual plantation map and the annual forest loss map.

### 2.4. Expansion of roads

Public roads are built and maintained by the government to facilitate the transport of goods and people from one settlement or city to another. The government of Indonesia defines its main public roads as national roads, provincial roads, and strategic roads. The Trans-Papua Highway and the Merauke Integrated Food and Energy Estate (MIFEE) highway are strategic roads made up of several national and provincial road segments, some still under construction, across Papua and West Papua, totalling > 4,000 km.

The annual expansion of main roads represents the length of road segment created each year. To map the expansion of main roads, we scanned available maps from local and national governments to identify their approximate routes. Next, we scanned the sequence of annual cloud-free images forward in time from 2000 to 2019 to detect road segments added each year along those approximate routes. We delineated road segments in a 1:50,000 scale using a visual, expert-based interpretation method. We used a professional Cintiq 22HD Wacom tablet with definition display and pressure-sensitive pen for the digitizing process. We did not distinguish paved versus unpaved road segments.

### 2.5. Other drivers of forest loss

We estimated the extent of forest loss caused by industrial-scale mechanized logging by mapping the annual expansion of all primary logging roads (large enough to be detected on Landsat images). We delineated logging road segments inside logging concessions using a similar digitizing procedure as we did for public roads. Next, we recoded forest areas underneath as ‘forest loss’ because the original Tree Loss dataset (v.17) is known to undercount losses caused by logging roads prior to 2011 (Hansen et al., 2013). Next, we defined a 700 m distance from logging roads beyond which timber extraction typically ends in Indonesia (Gaveau et al. 2014). We buffered logging roads by this distance and declared any forest loss within this buffer as forest loss caused by timber extraction. Inside known active mangrove logging concessions (Sillanpää et al. 2017) where clear-cutting (keyhole) timber extraction techniques are used instead of the typical logging road network, we recorded any loss of mangrove forest as natural timber extraction.

Inland Water body-related forest loss represents the area of forest that results from the expansion or shift of rivers and lakes between 2001 and 2019. To map these losses, we first created a detailed base map of inland water bodies (rivers and lakes) in 2000. We digitized small river lines at a 1:50,000 scale. We applied a supervised classification over the 2000 cloud-free Landsat composite to map larger rivers and lakes. Next, we combined our 2000 inland water bodies base map with a published map for changing water surface (Pekel et al., 2016), also developed with Landsat to measure the area where water expanded through surface run offs, expanding lakes, and changing river courses through to 2019. Within this area, we reclassified the annual cloud free composites (see §2.6) using a supervised classification to extract areas of forest converted to water instead of using the Tree Loss dataset (v.17) because we found that this latter dataset missed many of these events.

A burned forest is an area of forest characterized by deposits of charcoal and ash, removal of vegetation, and alteration of the vegetation structure. We estimated the areas of forest loss by fires by combining the forest loss map with active fire hotspots (NASA-FIRMS 2020). If an area of forest loss coincided with a cluster of fire hotspots, this area was recoded as: forest loss by fires.

We estimated the extent of forest loss caused by expanding population centres by measuring forest loss within 5 km of transmigration sites, regency and provincial capital cities and that did not qualify as any category of loss described earlier. We mapped transmigration sites using the 2000 cloud-free Landsat composite and government maps indicating their locations. We obtained the location of cities from government spatial data repository websites.

### 2.6. Annual cloud-free composites

We generated annual cloud-free Landsat composites for each year from 2000 to 2019 with Google Earth Engine (Gorelick et al. 2017). We masked cloud observations with the quality band ‘pixel_qa’, which is generated from the CFMASK algorithm and included in the Surface Reflectance products (Foga et al. 2017). Then, for each year, we created the annual composites using two criteria: 1) the median pixel-wise value of the Red, Near Infrared (NIR), and Shortwave Infrared (SWIR) bands of the images acquired between 1 January and 31 December, and 2) the minimum pixel-wise normalized burned ratio (NBR) of the images taken in the same given year. The composite image based on the median produces clean cloud-free mosaics but tends to omit new plantations and new roads developed at the end of the year. The second approach, based on the minimum NBR, produces noisier composites (residual clouds and shadows may persist), but it presents the advantage to capture plantations and roads that were developed at the end of the year.

### 2.7. Concession and protected area maps

Concessions are areas of land leased out to companies for the establishment and management of large monoculture plantations of oil palm or pulpwood (acacia), for mining activities, or for the extraction of natural timber. Pulpwood and logging (for natural timber extraction) concessions are regulated and issued only by the Ministry of Environment and Forestry. We downloaded pulpwood and logging concession datasets from government data repositories (MOEF, 2020). We downloaded the protected area dataset from the same data repository. Assembling the oil palm concession dataset was more difficult due to a range of ascending permits that are required by Indonesian law to earn the right to clear forest, plant and then manage oil palm plantations (Paoli et al. 2013). We assembled maps of oil palm concessions based on various datasets compiled by various Indonesian NGOs. Details are provided in supplementary methods in Appendix A.

### 2.8. Predictive modelling

We developed a spatially explicit model of deforestation (forest loss) using the principles of land-rent theory (Angelsen, 1999). In general, plantation companies and farmers act to maximize profits by allocating any parcel of land to the use that earns the highest rent. Topography (slope and elevation), roads, restricted zones (protected areas), and lands allocated for the development of plantations (concessions) determine the spatial distribution of land rent. Roads improve accessibility into remote forest areas and reduce transportation costs of agricultural commodities, which tends to promote deforestation. Steep slopes and high elevations limit accessibility, which tends to reduce deforestation. Accordingly, we built a set of six spatial predictors at a spatial resolution of 100 m × 100 m including slope and elevation, distance to main public roads expressed as a cost map, the presence of logging roads, protected areas and active concessions of oil palm, pulpwood and mining sites. We defined active concessions as those where we observed industrial activity between 2000 and 2019. We modelled distance to public roads (including the Trans-Papua and MIFE Highway) as a cost distance to simulate the propensity of people to travel along the path of least resistance. We calculated the cost distance as a function of slope as derived from the SRTM digital elevation model (Rabus et al. 2003).

We used weights of evidence and logistic regressions to generate deforestation-risk maps depicting the areas that are most at risk of deforestation. The risk maps have values ranging from 0 to 1, with higher values indicating greater vulnerability of deforestation. Weights coefficients were estimated for the categorical predictors (presence of active concessions, protected areas, and logging roads). Logistic regression coefficients were estimated for the continuous predictors (slope, elevation, and cost distance to public roads). The weights and logistic regression coefficients indicate the effect of predictors on forest loss. Positive weights/coefficients promote forest loss, negative ones reduce it, and close to zero values do not exert an effect. Since weights of evidence and logistic regressions assume that variables are spatially independent, correlation analysis between the predictor variables were performed. We tested several correlation methods. All tests highlighted no high correlation between paired variables. We performed the modelling using Dinamica Ego software (Soares-Filho et al. 2009).

We calibrated six models (each with different predictors) using the forest loss observed during the period 2010 – 2018 as the response variable. We chose this time frame because it captured most of variations in deforestation observed in the region. For each model, we generated yearly deforestation-risk maps, with probability values ranging from 0 to 1. To validate the model, we first simulated the spatial occurrence of forest loss from 2010 to 2018 using the average rate observed during this time frame. Simulation was performed via Cellular Automaton, in which the state of a cell is based on their current state and the states of the cells in its neighbourhood. Deforestation allocation and expansion are controlled by *Patcher* and *Expander* functions in Dinamica EGO. Next, we validated the simulated forest loss map against the map of forest loss observed during the same time frame (2010-2018) based on a vicinity-based comparison method implemented in Dinamica EGO. This method quantified the number of matching ‘deforested’ cells (one cell = 10,000 m^2^) between the observed and simulated maps and produced a similarity measure ranging from 0% to 100%. We selected the model that presented the simulation with the highest similarity index.

The selected model was used to predict forest loss over an 18-year time frame, i.e. to 2036. We developed two scenarios based on two hypothetical future speeds. In the first scenario, the speed of forest loss was estimated as the percentage of forest loss observed during 2010 – 2018 in Indonesian New Guinea relative to the total amount of forest left. In the second scenario, it was estimated as the percentage of forest loss observed in Indonesian Borneo during 2001 – 2018 (Gaveau et al. 2019).

## 3. Results

### 3.1. Situation in 2019

We estimate that 34.29 Million hectares (Mha), or 83% of Indonesian New Guinea was forest in 2019 (Figure 1). This “forest” comprises old-growth forests, including those impacted by selective timber harvesting (reclassified as “secondary” forest by the Indonesian Ministry of Forestry and Environment). This area of forest includes 30.30 Mha on drylands, and non-peat swamp habitats (‘other forests’), 2.82 Mha of peat-swamp forest and 1.17 Mha of mangrove forest. By comparison, the Ministry reported slightly less: 33.74 Mha of “primary and secondary” forest in 2017 (MOEF 2018). About one third of the non-forest area (6.86 Mha) was a savanna ecosystem, the Trans-Fly eco-region dominated by grasslands at the southern tip of the island in the Merauke Regency. Legally gazetted conservation areas, including reserves and national parks, protected 8.68 Mha, or 21% of the total land area.

**Figure 1.**
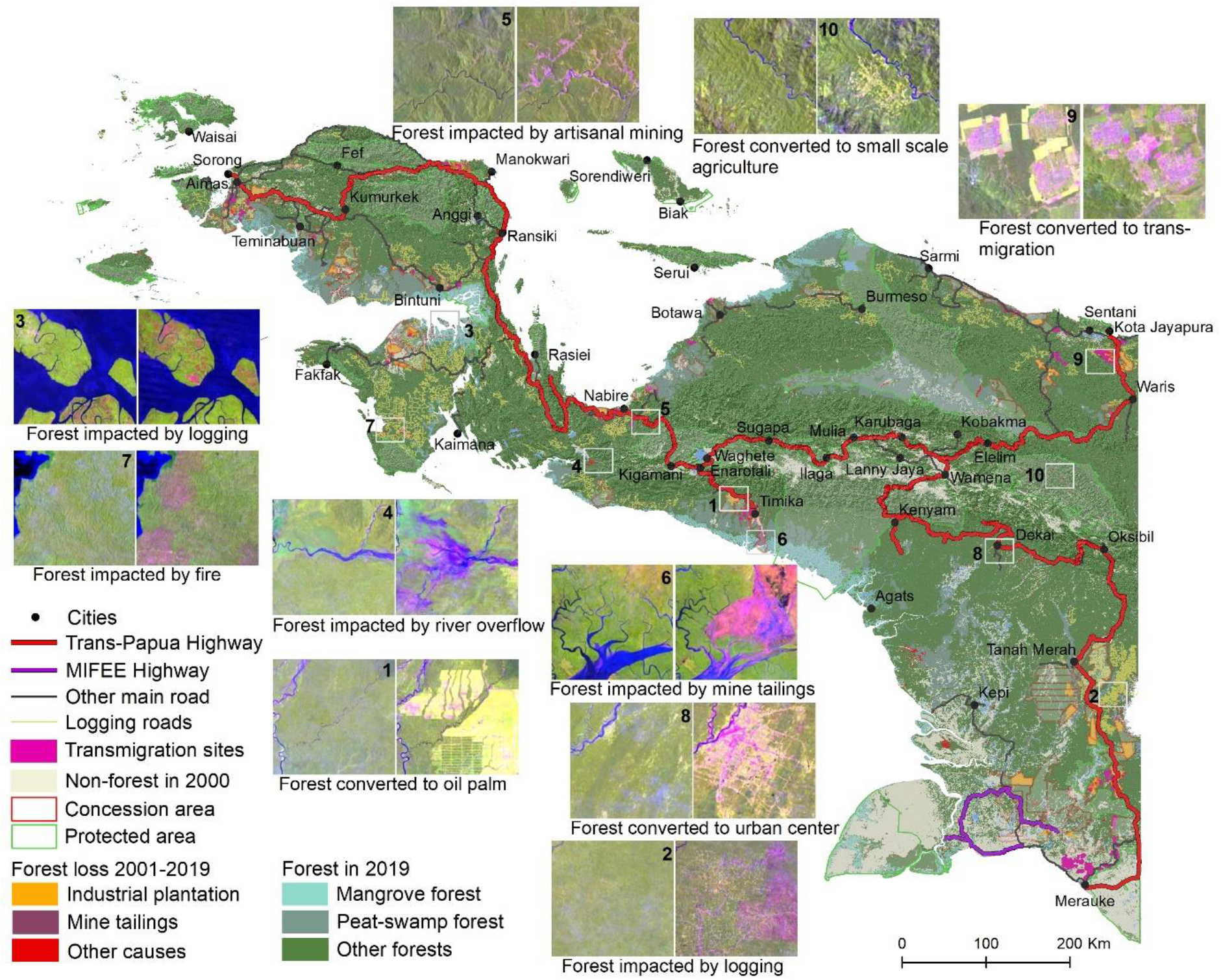
Land use and land cover change 2000 to 2019. All insets have the same scale.

We mapped 8,434 kilometres (km) of main public roads in 2019, including 3,887 km of Trans-Papua Highway (Red line; Figure 1), and 336 km of highway built specifically to develop the Merauke Integrated Food and Energy Estate (MIFEE) project in the south of the island (purple line; Figure 1). In addition, we mapped 22,035 km of wide (> 15 m) temporary trails built in the forest interior to extract natural timber (logging roads, Figure 1), translating into a density of logging trails of 0.064 km km^−2^ across the whole of Indonesian New Guinea.

We mapped 136 distinct industrial plantations (min = 85 ha; max = 16,882 ha; mean = 2,068 ha), covering a total planted area of 281,223 ha in 2019 (271,486 ha oil palm; 9,737 pulpwood). We found these plantations in 87 ‘active’ concessions from a total of 177 concessions given out by the government (1.81 Mha oil palm; 813,000 ha pulpwood). We also found one ‘active’ mining concession (198,000 ha) where mining company Freeport extracts gold and copper and where it dumps its wastes (tailings). Seventy four percent (2.10 Mha) of this total concession area (2.83 Mha) was still forest in 2019 (1.37 Mha oil palm; 577,000 ha pulpwood; 149,000 ha mining). Our planted area includes immature, damaged, and failed plantations and thus surpasses estimates that include only closed-canopy mature plantations (e.g., 175,803 ha, Descals et al. 2020). The Government of Indonesia’s estimates based on company reports (213,359 Mha) include mature and damaged plantations (DGOEC 2020).

### 3.2. Forest loss and drivers (2001 to 2019)

About two percent of old-growth forest, that is 748,640 ha, was lost between 2001 and 2019: 511,882 ha (1.9%) in Papua, 236,758 ha (2.6 %) in West Papua. we estimate that 45% (336,877 Mha) have become converted to plantations, mining, roads, and population centres, while 55% (411,763 Mha) have been impacted by transient processes such as selective natural timber extraction, inland water bodies-related processes, fires, shifting agriculture, cryptic timber harvesting and mining (Table 1). We break down these numbers by drivers below (see also Table 1).

**Table 1.**
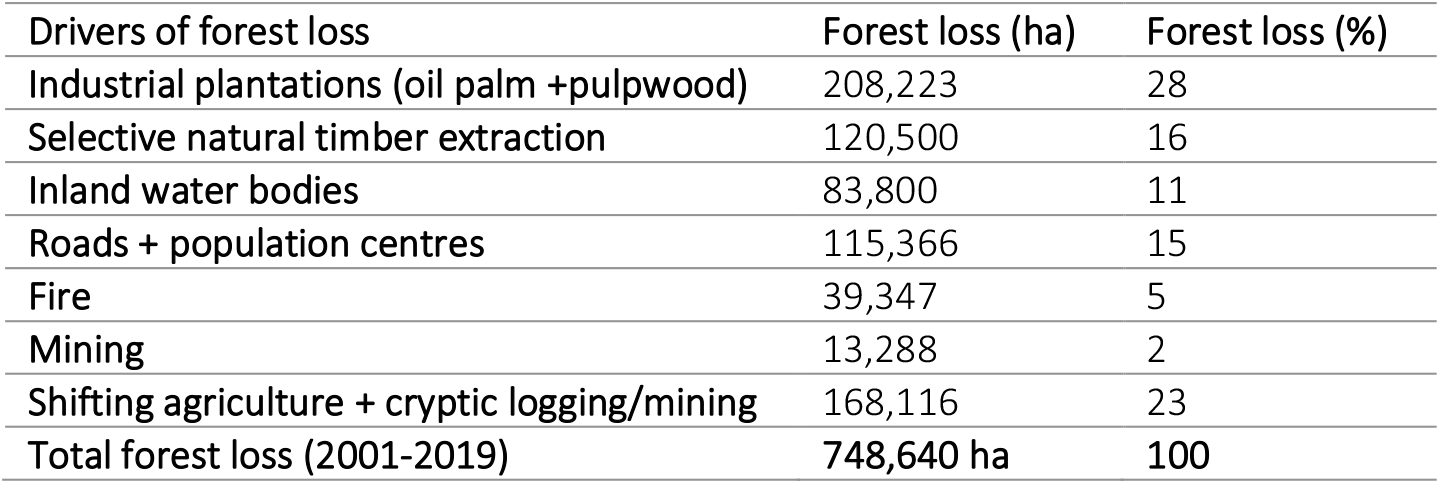
Summary of drivers of forest loss

We estimate that 208,223 ha of forest have been permanently converted to industrial oil palm and pulpwood plantations (97% oil palm; 3% pulpwood) since 2000, of which 97% were cleared and converted within the same year (203,202 ha; sum of all white bars in Figure 2). This conversion explains 28% of total forest loss. Oil palm and pulpwood concessionaires converted large rectangular blocks of old-growth forest to large oil palm and acacia estates (mean size = 2,068 ha) inside concessions. An example of this conversion is shown in inset 1 of Figure 1.

**Figure 2.**
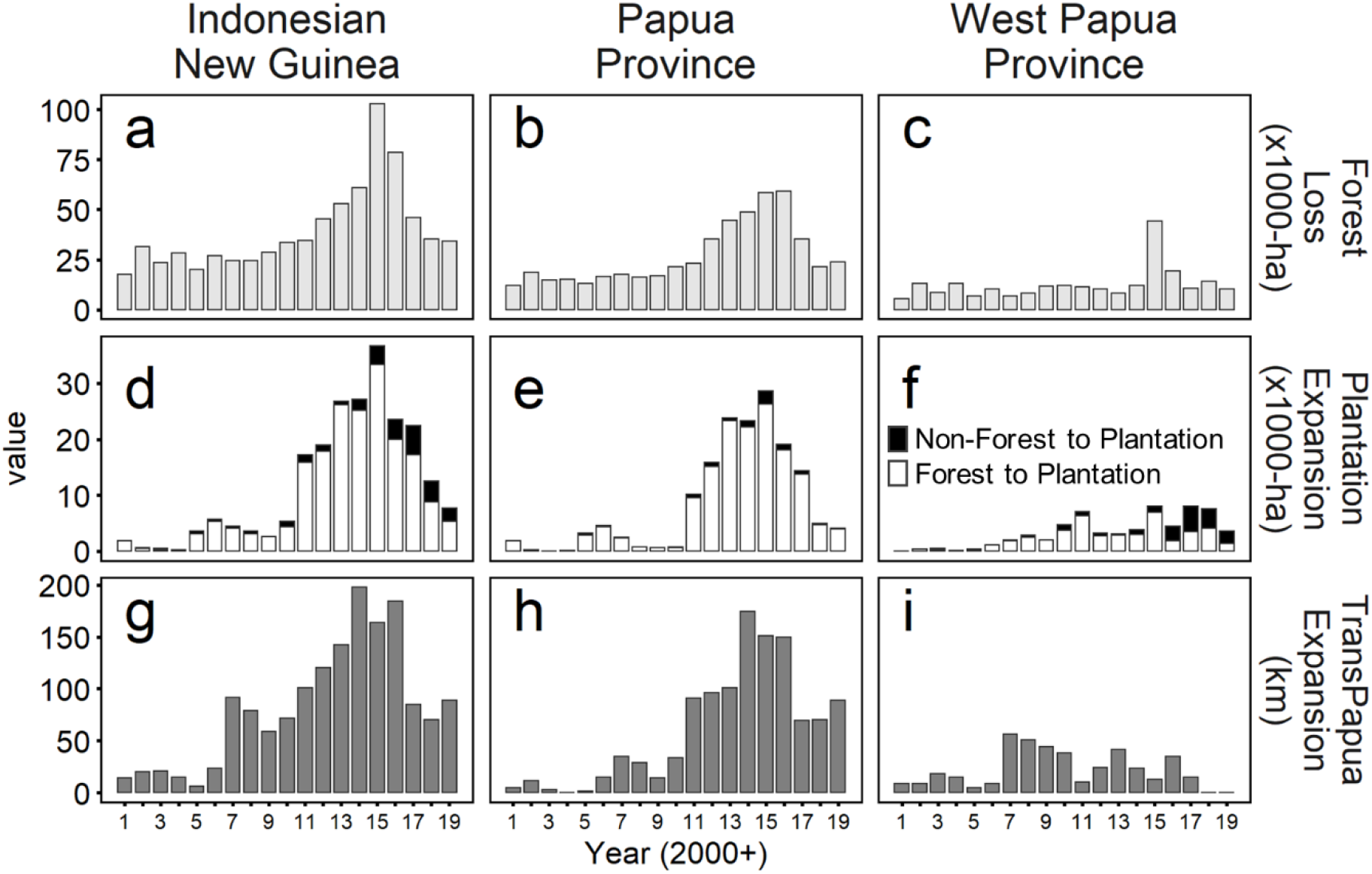
Indonesian New Guinea’s land-use change and its two provinces Papua and West Papua from interpretation of Landsat imagery year-by-year (2001-2009). Here we present the annual loss of forest area (a, b, c), the concomitant annual expansion of industrial plantations of oil palm and pulpwood (97% oil palm; 3% pulpwood) (d, e, f) and the expansion of the Trans-Papua Highway. The white bars in (d,e,f) represent the area of forest cleared and converted to industrial plantations in the same year.

We estimate that 120,500 ha (16% of total forest loss) were cleared by selective natural timber extraction. This estimate combines the loss of forest directly replaced by the 22,035 km network of extraction (logging) roads, by clearings near these roads (within 700 m) and by “keyhole logging” extraction patterns in the world’s largest mangrove concession (Sillanpää et al. 2017). This clearance is temporary, as the forest usually regenerates quickly along extraction sites (Gaveau et al. 2013; Sillanpää et al. 2017). Typical examples of natural timber extraction with logging roads and in mangrove forests seen by satellites are shown in insets 2 and 3 of Figure 1.

We estimate that 83,800 ha (11%) lost forest cover through inland water bodies-related processes including the expansion or shift of rivers and lakes. An example of this transient process is shown in inset 4 of Figure 1.

We identified 13,288 ha of forest cleared by mining activities (2%), including 1,000 ha of forest cleared by artisanal gold mining along the Trans-Papua Highway near the city of Nabire (see inset 5, Figure 1), and 11,372 ha cleared by tailings dumped in the Aikwa Delta system by Freeport’s Grasberg gold and copper mine located in the Sudirman Mountains in Mimika Regency (see inset 6, Figure 1). This result is corroborated by Alonzo et al. (2016).

We estimate, from fire hotspots and Landsat imagery, that fires impacted 39,347 ha (5%) of forest, including three large forest fires in October 2015 in West Papua (see inset 7, Figure 1). This clearance is likely temporary, as forest fires usually only burn the understory, and the vegetation regenerates after fire.

We determined that 115,336 ha (15%) were cleared by road expansion and population centres. This included forest directly cleared by the expanding route of public roads, and forest cleared within 5 km of transmigration sites, regency, and provincial cities. See for example, the rapidly expanding city of Dekai (inset 8, Figure 1) and expanding transmigration sites near the Jayapura city (inset 9).

Shifting cultivation and other processes, such as landslides, tornadoes, undetected understory fires, cryptic logging and mining activities are presumed to account for the remaining 166,920 ha (22%) that we have not characterised. An example of forest impacted shifting agriculture inside Pegunungan Jayawijaya Wildlife Reserve is shown in Inset 10 of Figure 1.

### 3.3. Deforestation trends

From 2001 to 2010 annual forest losses were relatively consistent, averaging 26,000 ha, before rising to 103,000 ha in 2015 and declining back reaching 35,000 ha in 2018 and finally 34,000 ha again in 2019 (Figure 2, panel a). These trends largely reflect Papua province, where forest loss peaked in 2016 (59,000 ha) and then reduced by more than half by 2019 (20,000 ha). In West Papua, trends were less pronounced, but also with a peak in 2015 (44,000 ha) when several large forest fires occurred in Fakfak Regency (see inset 7 in Figure 1).

### 3.4. Plantation expansion

By the end of 2000, there were 50,842 ha of industrial plantations (all oil palm). Between 2001 and 2019 the total area of plantations increased by 230,381 ha (220,644 ha oil palm; 9,737 ha pulpwood), with the majority (208,842 ha; or 28% of forest loss) replacing forests. Much of this expansion occurred in Papua Province (168,361 ha; 73%), and in the two southern regencies of Boven Digul and Merauke (109,585 ha; 48%), which contain just 15% of Indonesian New Guinea’s landmass. There appears to be three distinct periods for expansion. From 2001 to 2010 expansion was moderate and averaged 2,843 ha each year. In 2011, this expansion accelerated until it peaked in 2015 (38,000 ha added) (Figure 2d). In the third phase, expansion declined again though it remained higher than the average pre-2010 level. Plantations expansion from 2001 to 2019 focused on Papua Province (73%, 168,000 ha, Figure 2e). Forests converted to plantations within one year of forest clearance (white bars in panels d,e,f) exhibit similar trends.

### 3.5. Highway construction

By the end of 2000, 2,333 km of Trans-Papua Highway had already been built (construction started in 1979). Between 2001 and 2019 the Highway increased by 1,554 km, reaching 3,887 km in 2019. This included a 202 km road section built through Lorentz National Park (a World Heritage site), followed by 575 km added to link the remote cities of Kenyam, Dekai and Oksibil (Figure 1). In 2019, the Trans-Papua Highway was nearly complete though 51 km remained unopened. The construction of the Trans-Papua Highway generally increased from 2001 to 2016 (Figure 2g), from 14 km of road added in 2001 to 185 km in 2016, with a peak in 2014 (198 km added). In Papua Province (Figure 2h), there were two distinct periods: an average of 16 km added each year from 2001 to 2010 and then increasing to a maximum of 175 km in 2014. We see less variation in West Papua (Figure 2i). Roads expansion declined in 2017 (85 km) and 2018 (70 km), with little increase in 2019 (89 km). This drop reflects changes in Papua Province.

In Papua’s southern Merauke district, construction of the highway built for the Merauke Integrated Food and Energy Estate (MIFEE) began in 2012, totalling 336 km in 2019 (Purple line; Figure 1).

Transmigration population centres (totalling 129,427 ha, 90% established before 2000; Figure 1) and industrial plantations are all located within 35 km of these Highways.

### 3.6. Correlations

Variation in annual forest loss is positively correlated with the annual expansion of industrial plantations (r^2^=0.80, p<0.01; Figure 3a). This is clearer in Papua (r^2^=0.88, p<0.01) than in West Papua (r^2^=0.27, p=0.021) (Figure 3b&c). The annual expansion of the Trans-Papua Highway and plantations are positively correlated (r^2^=0.75, p<0.01; Figure 3). This relationship is strong in Papua but not in West Papua (r^2^=0.79, p=<0.01 versus r^2^=0.0088, p=0.70, Figure 3e,f). Similar patterns arise between annual highway expansion and forest loss (r^2^=0.67, p<0.01, r^2^=0.84, p<0.01, r^2^=0.19, p=0.57 for Indonesian New Guinea, Papua, and West Papua respectively) (Figure A2).

**Figure 3.**
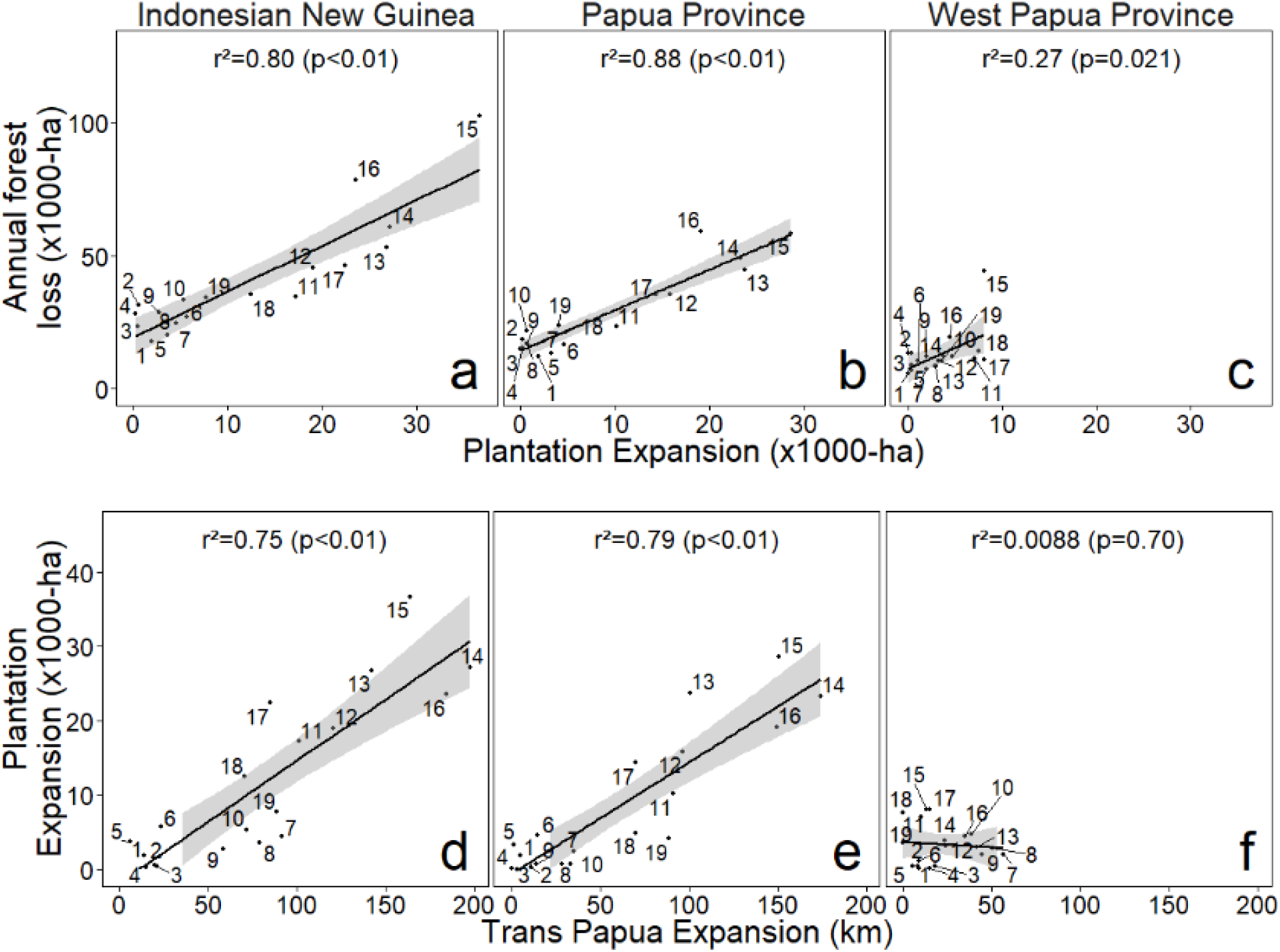
Scatter plots and associated correlation line and Pearson’s correlation coefficient (r^2^) between the annual expansion of plantations and the annual loss of forest (a, b, c) and between the annual expansion of the Trans-Papua road and the annual expansion of plantations (d, e, f) in Indonesian New Guinea (a, d) and its two provinces, Papua (b, e) and West Papua (c, f). The shaded areas show the 95% confidence interval for predictions from the linear model.

### 3.7. Predictive model

We calibrated six spatially explicit models of deforestation with the observed 2010-2018 forest loss (Table 2). We retained the model that included active concessions, protected areas, and slope-based road cost because it had the highest similarity (39% of observed forest loss were properly simulated) with the fewest parameters (Model 3 in Table 2).

**Table 2.**
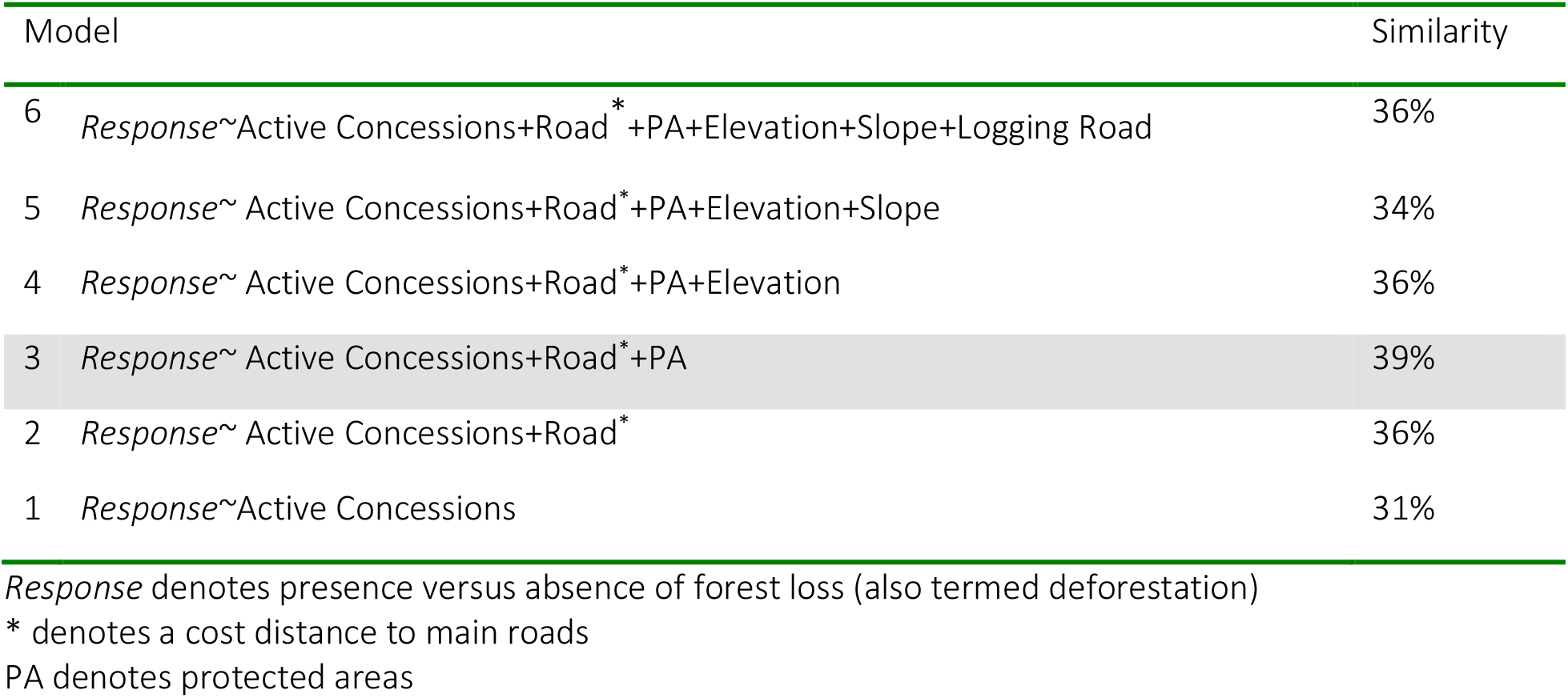
Spatially explicit models describing forest loss across Indonesian New Guinea.

| Model | Similarity |
| --- | --- |
| 6 *Response*~Active Concessions+Road*+PA+Elevation+Slope+Logging Road | 36% |
| 5 *Response*~ Active Concessions+Road*+PA+Elevation+Slope | 34% |
| 4 *Response*~ Active Concessions+Road*+PA+Elevation | 36% |
| 3 *Response*~ Active Concessions+Road*+PA | 39% |
| 2 *Response*~ Active Concessions+Road* | 36% |
| 1 *Response*~Active Concessions | 31% |

*Response* denotes presence versus absence of forest loss (also termed deforestation)
\* denotes a cost distance to main roads
PA denotes protected areas

We assessed the augmented threat posed by the observed expansion of roads by comparing the resulting deforestation-risk map against a hypothetical deforestation-risk map if roads had remained at their 2000 level. The construction of new roads increased the area of forest at risk of deforestation (probability > 0.66) by 30% compared to the hypothetical scenario of halting such developments. Lowland forests near the cities of Dekai and Kenyam in Papua province experienced the highest augmented threat (Figure 4). The forests near the cities of Tanah Merah Merauke, and Timika remained overall the most threatened. The threshold of 0.66 is the lowest probability value in which a forest pixel was seen converted to non-forest in our projections (Figure 5).

**Figure 4.**
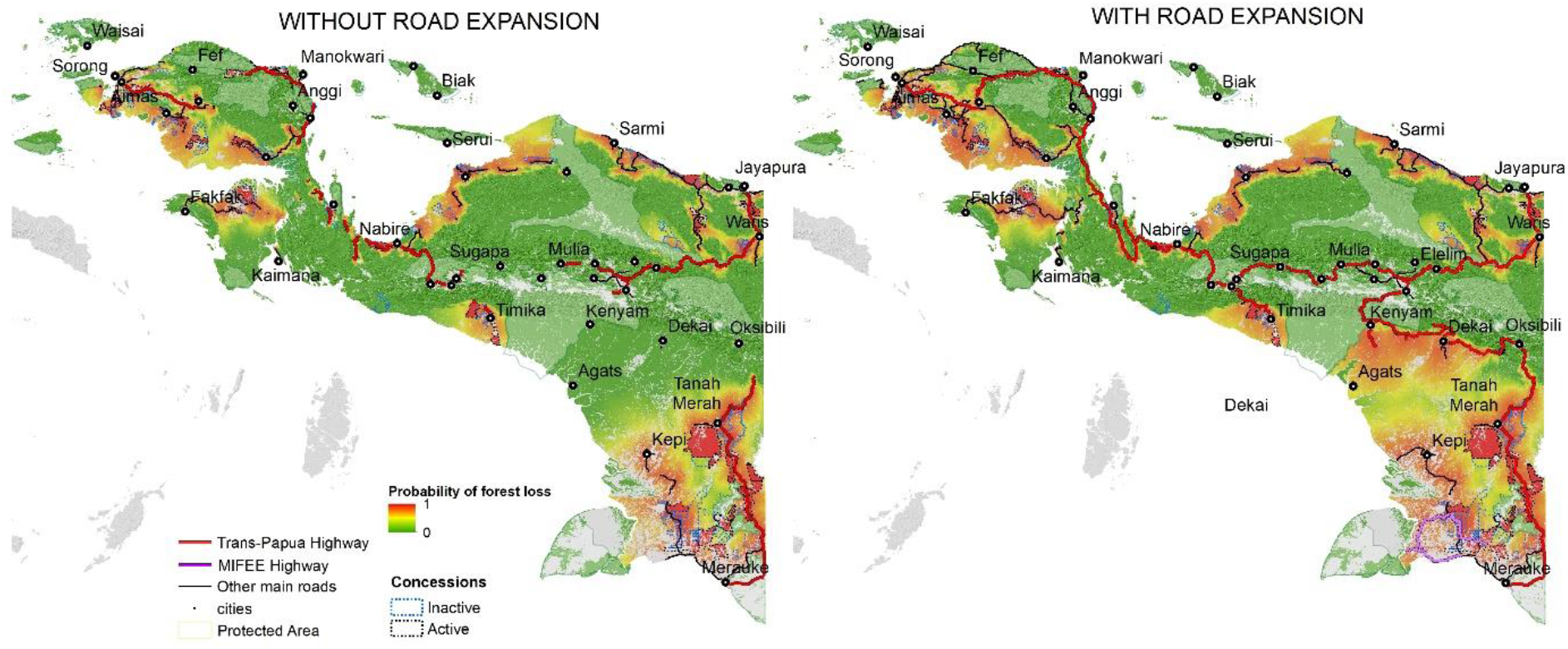
Map of Indonesian New Guinea showing a hypothetical deforestation-risk map without road expansion, i.e. as roads were in 2000 (Left Panel), and with the observed expansion of roads between 2001 and 2019 (Right Panel). Trans-Papua and MIFE Highways (red and purple lines) and other main roads (black). Areas of forest at high risk of deforestation are in red while low risk areas are green.

**Figure 5.**
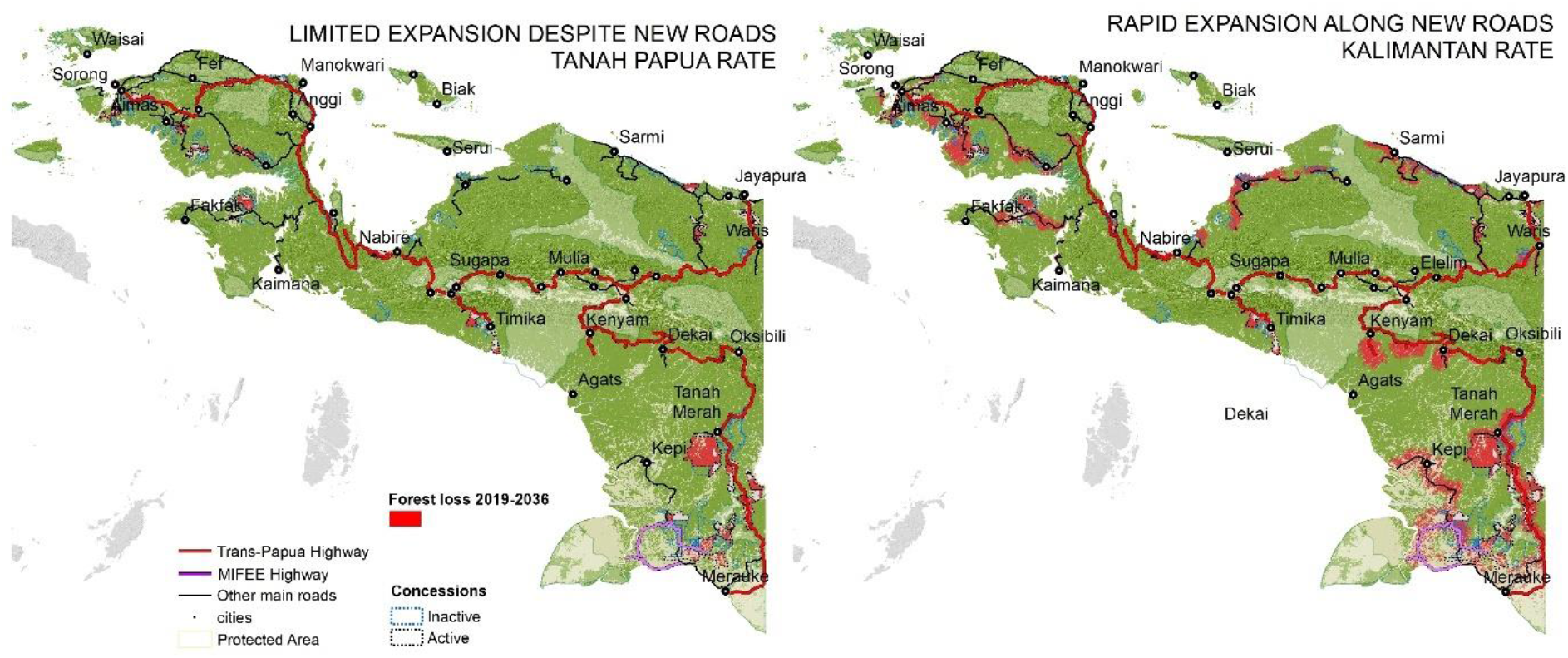
Forest conversion outlook to 2036 under two scenarios.

We projected future deforestation for 18 years, to 2036, for two scenarios using two empirically derived historic deforestation rates, one low and one high (Figure 5). In the first scenario, the speed of forest loss was estimated at around 55,843 ha yr^−1^ although we note that this rate varied slightly year on year since it was estimated as the percentage of forest loss during 2010 – 2018 in Indonesian New Guinea relative to the total amount of forest left in the preceding year. This is the ‘Limited Expansion Scenario’ despite new roads. In the second scenario, using the speed of forest loss experienced by Indonesian Borneo during 2001 – 2018, we estimated forest loss at 253,907 ha yr^−1^. This is the Business-as-usual scenario of rapid expansion seen in Indonesian Borneo following roads construction, and strong government support for plantation expansion.

In the limited expansion scenario, a total of 932,119 ha of forest would be lost by 2036, with 701,335 ha alone in active concessions (596,715 ha oil palm; 93,974 ha pulpwood; 10,646 ha mine tailings), 35,636 ha in inactive concessions, and 195,148 ha along roads. In the business-as-usual scenario a total of 4.50 Mha would be lost by 2036, with 720,353 ha in active concessions, 430,147 ha in inactive concessions, and 3.2 Mha along roads. The forest between the cities of Kenyam and Dekai, and between Tanah Merah and Merauke, would be the most severely impacted (Figure 5).

## 4. Discussion

We have mapped and described trends in forest loss, industrial plantations and roads expansion, and their overlap in Indonesian New Guinea each calendar year from 2001 to 2019. We showed that the presence of concessions, roads and protected areas influenced the spatial distribution of forest loss and used them to predict future losses under different scenarios. These datasets are presented interactively in the Papua Atlas (Papua Atlas 2021), a platform developed to monitor the impacts of expanding roads, mines and industrial plantations on forests, and thus assist the provinces oversee their commitments under the Manokwari Declaration (Cámara-Leret et al. 2019). Overall, these spatial data as explored in this study, provide information for spatial planning across this important, yet understudied region.

About 34.29 million hectares (Mha) of forest remained in Indonesian New Guinea in 2019, representing 42% of Indonesia’s forests. This is Asia Pacific’s largest area of intact old-growth forest. Furthermore, we find that Indonesian New Guinea has 8.5% (1.17 Mha) of the World’s mangrove forests (13.58 Mha), with Papua’s Mimika coast and West Papua’s Bintuni Bay home to Indonesia’s largest contiguous mangrove blocks (340,000 ha and 222,000 ha), only second to the Sundarbans in Bangladesh (Bunting et al. 2018; Global Mangrove Watch 2021). Given their importance to fisheries and role in carbon sequestration, conserving these mangroves is crucial (Murdiyarso et al. 2015). Similarly, we find that with 2.82 Mha of intact forest occurring on “officially recognised” peat, Papua and West Papua provinces have Indonesia’s largest remaining intact peat forests. There are large uncertainties in these estimates (Warren et al. 2017) and recent ground truthing by Bappenas have highlighted a need to revise the current peat maps for Indonesian New Guinea (Imam Basuki pers. comm. to DS 2021).

What are the specific conservation impacts of the changes we observed? Our knowledge of the region’s biodiversity is sufficient to underline that many unique taxa will be at increased risk of loss as old growth forests are converted and loss. In line with patterns seen elsewhere these risks will be further compounded by forest fragmentation, degradation, and increased accessibility (Maxwell et al. 2016). Nonetheless, our spatial knowledge of these sometimes local and vulnerable species remains limited (Cámara-Leret et al. 2020; Marshall and Beehler, 2007; van Heist et al. 2010). We have not attempted any formal spatial priority setting here, though this is much needed and would be a valuable complement to our analysis. Such an exercise would reflect the new information generated over the last two decades (e.g., Volleringet al. 2016, Cámara-Leret et al. 2020) but will also be vital to highlight the many remaining unknowns (see e.g. Supriatna et al. 1999).

Overall, Indonesian New Guinea has suffered less forest loss and degradation than much of Indonesia. Just two percent of old growth forests were cleared from 2001 to 2019 and we estimate that more than half of this will recover if it is not further disturbed or converted. The density of primary logging roads (0.064 km/km^2^) was lower, around one seventh, what we had seen previously in Borneo (where using similar road mapping techniques, Gaveau et al. (2014), found 0.48 km/km^2^) and similar to the densities observed in Central Africa during the 1980s and 1990s (Laporte et al. 2007). Nonetheless, extensive forest cover means that most developments, impact these unique ecosystems. We showed that 88% of new industrial plantations (oil palm and pulpwood) replaced forests and accounted for 28% of observed deforestation. Mining remains a concern too. Notably, for over three decades the Grasberg Mine, the World’s largest goldmine and second largest copper mine, has been releasing approximately 230,000 tons/day of tailings into rivers that flow into the Aikwa Delta in Mimika Regency (Taberima et al. 2020). While we show that the total forest loss (11,372 ha) resulting from this process contributes just 2% of total forest loss across the Indonesian New Guinea, nonetheless the ecological and human impacts are considerable (Perlez and Bonner 2005).

Road developments have played a fundamental role in determining the pace and distribution of forest loss. The national government has invested heavily in roads, with the Trans-Papua Highway and the Merauke Food and Energy Estate Road intended to boost expansion of plantations and food estates. Our analysis clarifies the pace and impact of these developments. Trends in construction (annual number of kilometres created) of the Trans-Papua Highway are correlated with annual expansion (number of hectares added) of industrial plantations, indicating the synergy between road and plantations expansion. We observed a surge in activity after 2010, peaking at 2014-2016 while in other regions of Indonesia the expansion of plantations had slowed several years earlier as prices of crude palm oil declined and the global economy slowed (Gaveau et al. Under review). This recent peak is particularly marked in Papua province, which continued to attract new plantations for several years after expansion in other provinces had declined. We note that the observed correlation between roads and plantations need not be inevitable. While plantation developments require access, with sufficient controls in place new roads need not imply new plantations—though it is unclear if sufficient controls can be mustered in Indonesian New Guinea to permit such developments. Here we will not attempt a comprehensive review of the possible costs and benefits of roads as these have been reviewed many times (Laurance et al. 2014; Laurance and Arrea 2017). Nonetheless we note that the financial costs are often greater than anticipated. Projects such as the Trans-Papuan Highway that cross rugged rain-soaked terrain are challenging to develop and maintain due to landslides, collapse and floods (Alamgir et al. 2017). Whatever their engineering requirements new roads bring developments that may be viewed differently depending on perspectives. UNESCO has noted that the threats posed by new roads to the globally recognised values of the Lorentz National Park, a World Heritage Site, may require it to be designated as a World Heritage Site ‘in danger’ (Sloan et al. 2019). In addition to such recognised high-profile concerns, we see many other less known issues. For example, our observations show that the Trans-Papua Highway has already attracted artisanal gold mining in Nabire Regency, massive expansion of industrial plantations in Merauke and Boven Digul, and the rapid growth of Kenyam and Dekai Towns since they have been linked by 575 km of Trans-Papua Highway opening these areas to external market forces. While roads and associated development can bring welcome benefits to previously remote and neglected communities this requires detailed planning and is often debateable in practice. At least one recent study contends that the road infrastructures being developed in West Papua province’s Tambrauw regency will primarily benefit companies rather than local communities (FWI 2020). According to the study, neither past nor planned road construction in the regency were designed to help people access health care or other services or opportunities. Instead, they facilitate access by concession owners to their oil concessions and benefit their subsequent extractive industries with consequent harm to forests, to indigenous peoples and to biodiversity (FWI 2020). Concern that commercial interests dominate over local needs, remain common across the region (Boissière et al. 2013; Ariutama and Fahmi, 2019). We note that the Merauke Integrated Food and Energy Estate (MIFEE) launched in 2011 to develop over 1.2 million ha of rice, soya, sugar, cassava and oil palm, and triggered the construction of the MIFEE Highway described in this study as well as concession permits granted for industrial oil palm and pulpwood (mainly Acacia) plantations have displaced numerous indigenous Marind-Anim communities (Chao 2019).

The Minister of Environment and Forestry issued a regulation in 2020 that fast tracks permits previously protected forest(*Hutan Lindung*) to be converted for food production (Jong 2020). Such policy changes favour plantation developments at the expense of forests and the careful planning needed to account for the needs and concerns of indigenous communities. In the latter part of 2020, partly in response to COVID 19, the Indonesian national government already announced an expansion of the food estate model to include Merauke Mappi and Boven Digul Regencies, ostensibly to reduce dependence on food imports. Cheaper transport will reduce the cost of basic commodities including cement and other construction materials, bringing developments and new opportunities for plantations, mining, and other developments in many forest rich areas. Our model indicates the likely nature, magnitude, and location of future threats to the region’s forest. If development follows a top-down “business-as-usual” model similar to what has already occurred in Indonesian Borneo, our model suggests that an estimated 4.54 Mha of additional old growth forest could be lost by 2036. The relatively pristine forest regions surrounding Kenyam and Dekai, and in Boven Digul, Mappi and Merauke regencies comprising many important ancestral lands of indigenous groups are likely to experience significant deforestation if new roads bring the anticipated plantations, mining, food estates and associated influx of labour. Is there an alternative development scenario that will preserve the rich forest and cultural heritage of Indonesian new Guinea? First it is helpful to look briefly at the regulatory context.

The history of Indonesia’s state interventions concerning forests is complex but key aspects provide context. Indonesia’s constitution recognizes traditional ownership (*hak asal-usul*, Article 18) over forest but also declares, in an apparent conflict with traditional rights, that it is the state that has responsibility for controlling the use of the nation’s natural resources (*menguasai*, Article 33) (Lynch and Harwell 2002; Wrangham 2002). This contradictory message permitted post-independence governments to assert ever greater control over forests through a series of laws and regulations (Wrangham 2002). However, the constitution remained unchanged, and Indonesia’s Constitutional Court has ruled that customary forest lands (*hutan adat*) cannot also be State Forest land and that the government’s past claims to vast areas of forests were and are invalid (Butt 2014). Thus this ruling (No. 35 of 2012) and various subsequent laws have recognised customary claims to ancestral territories (Larson et al. 2016). The central government’s response has been high-profile commitments to improve the legal rights of Indonesia’s indigenous people and to return many forests to local communities. This process has raised various concerns, and there have been various efforts to re-impose government controls and oversight.

Within this history of contested claims several initiatives to reconcile forest protection and local needs have been developed around government led “Social Forestry” programs. Within these initiatives the government maintains overall authority and the emphasis is on communities being granted rights to manage and draw value from forest under a set of rules, rights, and obligations. It is in the context of such efforts combined with increased recognition of customary lands that many authors propose and promote some form of “social forestry” (for example, sago, candlenut and nutmeg production) as an alternative to industrial-scale oil palm, pulpwood and food estates (e.g., Fatem 2019; Ungirwalu et al. 2019). Nonetheless, such government controlled social forestry programs have had mixed success (Fisher et al. 2019); different land use contexts required different investment strategies to ensure social and environmental benefits (Meijaard et al. 2020). Whatever the outcome, social forestry appears to be the model of choice by the provincial governments of Papua and West Papua (see Manokwari Declaration) for engaging with customary communities on their terms (Fatem 2019). However, the status of customary forests remains unclear due to weak political will and incomplete procedures for recognition (Myers et al. 2017). As a result, the recognition of customary forests and awarding of social forestry permits by government authorities, have been slow (Meijaard et al. 2020).

There are more radical approaches to land-use planning and implementation that recognise the primacy of indigenous rights and attempt to reconcile multiple, sometimes competing, demands (Wollenberg et al., 2009; Boissiere et al., 2010; Sarkar et al., 2017). Given the long history of local control over their own territories along with the identification oversight and protection of their natural resources, these approaches appear well suited to Indonesian New Guinea. There is a need to integrate development and environmental constraints, with local preferences in a manner that indigenous people can benefit from and thus favour and support, and that can also include explicit conservation goals to the degree that these are acceptable to stakeholders (Wollenberg et al., 2009; Boissiere et al., 2010; Padmanaba et al., 2012). Relevant evidence that such approaches have value for both conservation and people is available from local examples from within Indonesian New Guinea showing that—at least in remote regions where government oversight has been largely absent—community-based oversight has been effective in protecting large areas of land and forest in a near pristine state (Sheil et al. 2015). In addition, there is often strong support from traditional communities in Indonesian New Guinea for land-use planning, with official recognition, that designates large areas, as identified by the communities themselves, to be protected in perpetuity (Sheil and Boissière 2006; Boissiere et al., 2010; Padmanaba et al., 2012; Sheil et al. 2015; Van Heist et al. 2015).

In any case, we favour approaches that recognise that indigenous communities have clear rights to their lands and resources that do not depend on anyone else’s permission. Ensuring such rights appears consistent with Indonesia’s constitution and would be particularly valuable in Indonesian New Guinea. Land-use planning in Papua needs to be based on a solid understanding of the benefits of economic development versus its social and environmental costs (Boissiere et al. 2010; Padmanaba et al. 2012). Papuans revere their customary practices and institutions making these a strong basis for building conservation and development opportunities around what local people themselves find important and desirable (Sheil and Boissière, 2006; Sheil and Lawrence, 2004; Van Heist et al. 2015). These local practices, though often labelled “traditional”, remain flexible and will adapt to changing requirements but can nonetheless benefit from external recognition and support when addressing novel threats, especially those that arise externally (Sheil et al. 2015). Strengthening this bottom-up decision-making process requires avoiding outdated top-down approaches. Effective, consensual, and frequent engagement between local representation and all layers of government is key to ensure alignment of local, regional and national development planning.

## 5. Conclusion

Indonesian New Guinea is opening to large-scale development. While overall losses remain relatively localised, there are significant local impacts and a high likelihood of more. Our models show how ongoing road development may lead to more extensive deforestation. Future land use decisions should carefully weigh the positive and negative impacts of developments within this ethnically and biologically diverse region. Listening to, and seeking guidance from, the people impacted by these decisions is crucial for avoiding harmful developments. This is a global concern, and we call on the international community to support the Indonesian and Papuan people and their elected governments in making the best decisions.

## 6. Appendix A

Contains Supplementary figures A1 and A2 and supplementary information describing how we created the oil palm concessions dataset.

## 7. Acknowledgements

This work was partly funded by DFID Indonesia, UK Climate Change Unit in collaboration with CIFOR and by the Rainforest Foundation Norway (RFN). DS’s time was covered by NMBU. The authors would like to thank Chip Fay from RFN for providing oversight and for his valuable comments and discussions throughout this project.

