## Supplementary material for "Forest loss in Indonesian New Guinea: trends, drivers, and outlook": Figure A1, Figure A2, Supplementary methods

**Appendix**

**Supplementary figures**

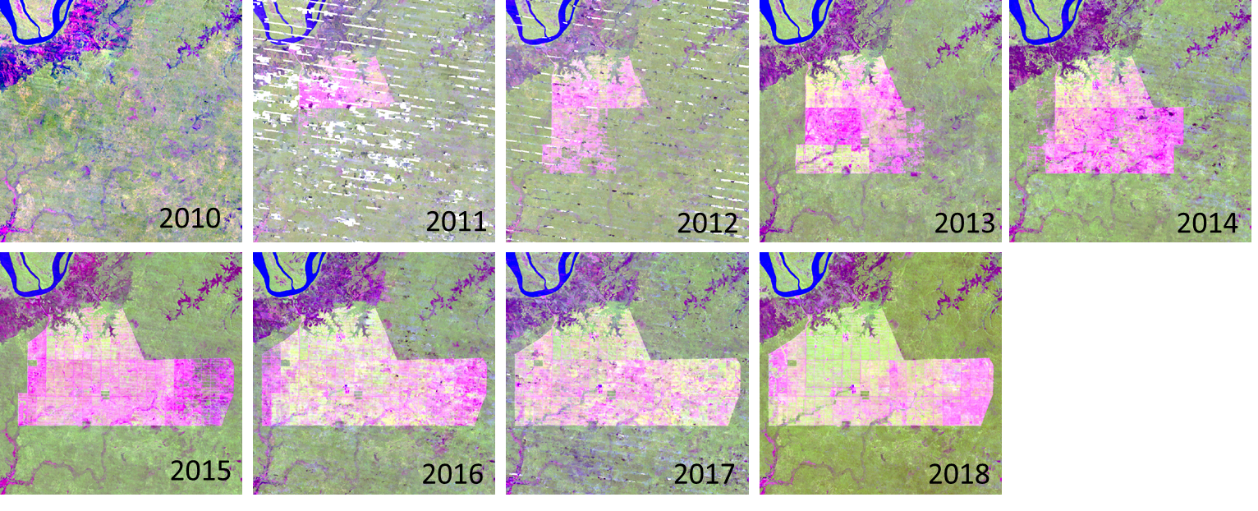

**Figure A1.** A sequence of cloud-free annual composite LANDSAT image snapshots revealing the annual expansion of an industrial plantation in Papua province, Indonesian New Guinea. Imagery displayed in false colors (RGB: Short-wave infrared: band 5; Near infrared: band 6; Red: band 4). Here, forest appears green, while recently cleared areas appear pink.

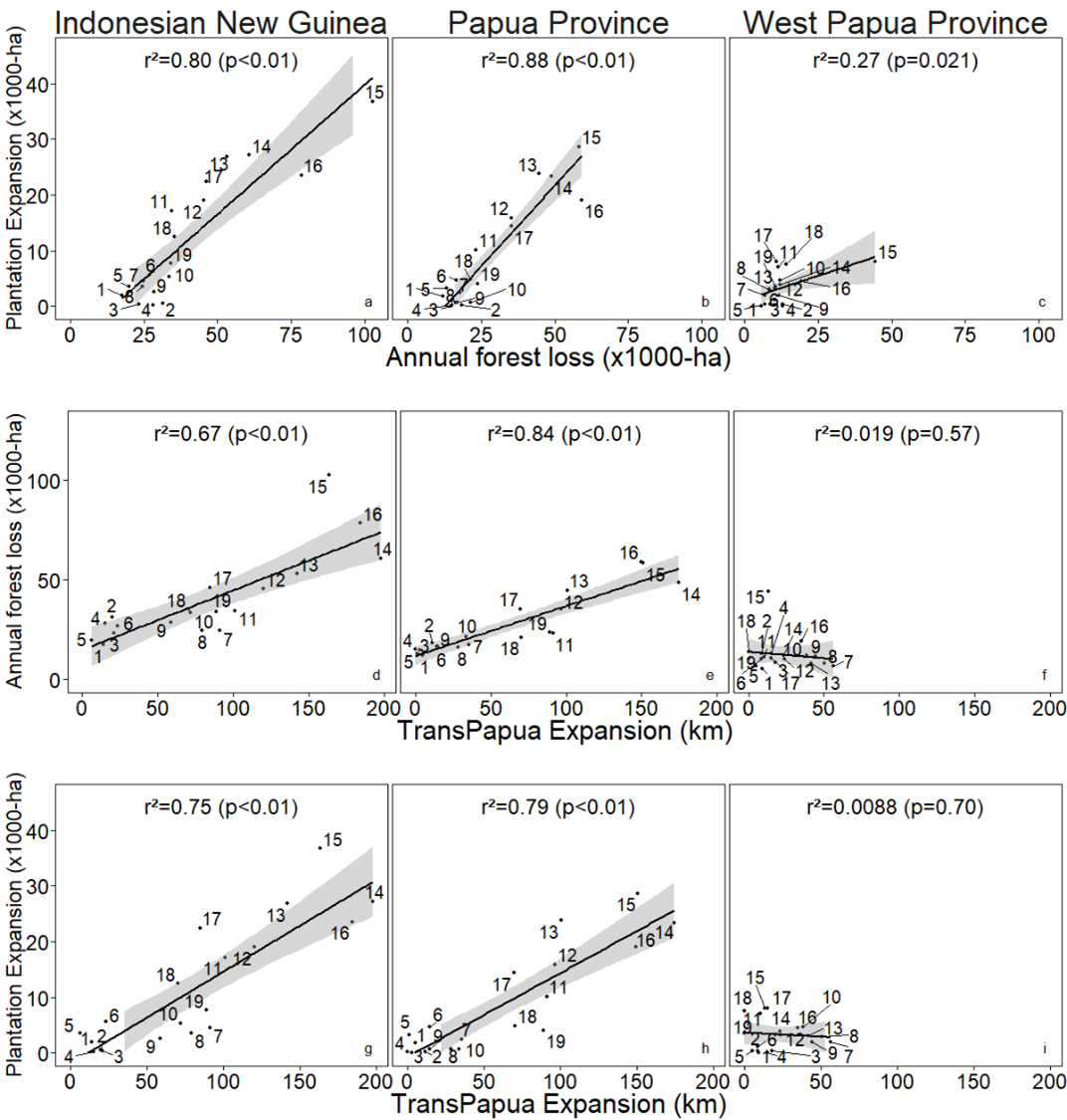

**Figure A2.** Scatter plots and associated correlation line and Pearson's correlation coefficient (r^2^) between the annual expansion of plantations and the annual loss of forest (top graphs), between the annual expansion of the Trans Papua road and annual forest loss (middle graphs), and between the annual expansion of the Trans Papua road and the annual expansion of plantations (bottom graphs) in Indonesian New Guinea and its two provinces, Papua and West Papua. The shaded areas show the 95% confidence interval for predictions from the linear model.

**Supplementary Methods**

**Oil Palm concessions**

Oil palm permits are issued by different levels of authorities (different ministries, their line agencies at district and provincial level, and district heads). As a result, the full information is scattered across different government institutions, and often maintained without a uniformed format. We assembled maps of oil palm concessions based on various datasets compiled by Indonesian NGOs, and online documents^[1]^.

The first significant step of an oil palm concession leasing process is to apply for a Location Permit (*Izin Lokasi*, ILOK). The company makes an application to the district head (the regent). If it is successful, the district head issues the permit giving the company the right to negotiate to acquire land within a given area, with local communities, and with the Ministry of Forest and Environment. The second step is the Environmental Impact Assessment, evaluated by the provincial environmental agency. If it is approved, the district head issues and environmental permit. The third step is the Plantation Business Permit (*Izin Usaha Perkebunan*, IUP), which grants the company the right to operate on land and is delivered by the district head or the governor. If the planned concession falls within State Forest area (*Kawasan Hutan*), it must be released by the Ministry of Forestry and Environment prior to forest clearing. Companies must apply to the Minister of Forestry, who may or may not relinquish the Ministry’s claim to the land by issuing a decree of State Forest Release (*Pelepasan Kawasan Hutan*, PKH). The final permit is the Land use right (*Hak Guna Usaha*, HGU). It is issued by National Land Agency (BPN) under the Ministry of Agrarian and Spatial Planning. It rounds off the permit process giving the company the tenure within a given boundary of land for 35 years. This permit can be extended.

After combining all available oil palm concession datasets, we defined a new dataset that took the maximum concession area extent by merging the boundaries of all permits for each concession. For example, if a third of a concession area had a HGU permit, another third had a PKH permit, and another third had an ILOK/IUP permit, the final concession extent used in this study was the sum of all three areas.

1. Datasets developed by Auriga, Greenpeace and Daemeter consulting. Online reports from Awas Mife, Pusaka, Gecko project
